# Distinct Roles of Surface Nanostructure and Polymer Degradation in Fibroblast Response to Device Design

**DOI:** 10.1101/2025.10.24.684439

**Authors:** Kendell M Pawelec, Erik M. Shapiro

**Affiliations:** Department of Radiology, Michigan State University, East Lansing, MI 48824, USA; Institute for Quantitative Health Sciences and Engineering, Michigan State University, East Lansing, MI 48824, USA; Department of Biomedical Engineering, Michigan State University, East Lansing, MI 48824, USA; Department of Chemical Engineering and Materials Science, Michigan State University, East Lansing, MI 48824, USA; Department of Physiology, Michigan State University, East Lansing, MI 48824, USA

**Author notes:** Corresponding author post-publication: Erik M. Shapiro.

**Keywords:** fibrosis, nanoparticles, degradation, foreign body reaction, polymer devices

## Abstract

Fibrotic encapsulation of implanted biomedical devices represents the highest cause of device failure across numerous fields, including orthopedics, breast implants and neural electrodes. While most research has focused on the immune system for driving the foreign body response to devices, fibroblasts are emerging as another key regulator of tissue response. In the present study, we evaluated the effects of porous device design on the activation of primary human dermal fibroblasts into myofibroblasts, a contractile cell that is a hallmark of fibrosis, over four weeks in culture. In particular two factors were studied 1) the introduction of nanoparticles, as they are increasingly being used for device functionalization, and 2) the polymer matrix, which dictates chemistry, mechanics and degradation. The addition of tantalum oxide (TaOx) nanoparticles (0-20wt%) had a minimal effect on fibroblasts, significantly down-regulating expression of alpha smooth muscle actin (αSMA) (0.31 ± 0.1), likely driven by the increase in nanoscale surface roughness. The polymer matrix of the device caused significant changes to myofibroblast activation, with the fast degrading polymer, poly(lactide co-glycolide) (PLGA) 50:50, significantly up-regulating multiple myofibroblast markers, including αSMA (5.16 ± 2.5), vinculin, integrin β_1_, and integrin β_5_, compared to non-degrading polycaprolactone (PCL). The effect was due to the release of degradation products, namely lactic acid, which affects cellular metabolism. Together this highlights that device design affects biological response immediately post-implantation in ways that impact the ultimate success or failure of biomedical devices.

## 1. Introduction

The biological response to implanted biomedical devices, known as the foreign body response, is a highly orchestrated interplay between device and cells, involving proteins and cytokines [1]. This complex process begins immediately post-implantation as proteins absorb onto device surfaces, subsequently triggering an immune response [1,2]. What follows are a series of steps including the deposition of extra-cellular matrix (ECM) and the activation of fibroblasts and other mesenchymal cells into myofibroblasts, that exert contractile forces on the ECM to close wounds [3]. With the eventual maturation and remodeling of surrounding tissues to achieve normal function, the foreign body response is analogous to normal wound healing [4]. However, when acute inflammation and myofibroblast activity persist as a chronic response, biomedical devices can become encapsulated, or coated in a thick layer of ECM, that is a hallmark of pathological wound healing and fibrosis [2].

The triggers of fibrotic encapsulation around implanted devices are not well understood. However, encapsulation exerts a devastating impact on device performance that can ultimately lead to failure across a spectrum of devices [2]. In orthopedics and breast implant surgery, encapsulation of devices contributes to movement of the device, pain and ultimately failure [5,6]. Where devices function as intraneural electrodes or sensors, a fibrotic tissue layer disrupts electrical signals, and therefore patient outcomes [7]. In all cases, failure due to fibrotic encapsulation is prevalent, and requires revision surgeries that increase the economic and societal burden of healthcare.

Given the high incidence of peri-implant fibrosis, it is important to understand which factors drive device encapsulation. In many cases, studies have focused on inflammatory processes as the key driver of fibrosis, since neutrophils, monocytes and macrophages are the first cell types to react to foreign materials [1]. Increasing evidence has shown a role for fibroblasts in the foreign body reaction, as they have an ability to exchange information with the immune system and modulate local and systemic responses [4]. Fibroblasts are also a major contributor to the myofibroblast population that is a hallmark of fibrotic diseases [3].

The materials properties of biomedical devices, including stiffness and surface topography, are known to affect both the immune response and severity of fibrosis [2]. We hypothesized that nanoparticle incorporation, used increasingly to fine tune device functionality, and polymer degradation products would play a significant role on myofibroblast activation. To test this, porous polymer matrices mimicking biomedical devices were cultured in the presence of human dermal fibroblasts. Activation of myofibroblasts was quantified after the incorporation of 0-20wt% nanoparticles into a polymer matrix, and as a function of the polymer degradation products released from non-degrading, slow degrading or fast degrading polyester matrices. It was found that nanoparticle addition decreased myofibroblast activation, while polymer degradation products could promote increased activation via metabolic modulation. Together this offers insight into the complexity of the foreign body response and considerations when designing the next generation of biomedical devices.

## 2. Experimental

### 2.1 Materials

Unless noted, all materials were purchased from Fisher Scientific. The polymer matrices used were polycaprolactone (PCL), poly(lactide-co-glycolide) (PLGA) 50:50 and PLGA 85:15. PCL (Sigma Aldrich) had a molecular weight average of 80 kDa. PLGA 50:50 (Lactel/Evonik B6010-4) and PLGA 85:15 (Expansorb® DLG 85-7E, Merck) were both ester terminated and had a weight average molecular weight of 80-90 kDa. All three are biocompatible polymers used in clinically approved implants.

### 2.2 Tantalum Oxide (TaO_x_) nanoparticles

Hydrophobic TaO_x_ nanoparticles (3-9nm diameter) were manufactured based on our previous procedure with one change [8]. Hydrophobicity is introduced by coating nanoparticles with hexadecyltriethoxysilane (HDTES, Gelest Inc., cat no SIH5922.0), an aliphatic organosilane. The coating has been shown to produce polymer composites with homogeneous nanoparticle distributions [9].

### 2.3 Composite manufacture: devices and films

Porous devices emulating tissue engineering implants were used for 3D cell culture and polymer films, mimicking the device walls, were used to study the effects of degradation products, as described previously [10]. For devices, polymers were solubilized in suspensions containing TaO_x_ nanoparticles (where applicable) in dichloromethane (DCM, Sigma); PCL was used at 8 wt %, while PLGA was used at 12 wt%. The amount of TaO_x_ nanoparticles (0-20wt%) was calculated based on the weight percent of the total dry mass (polymer + nanoparticle). To introduce microporosity to the device walls, sucrose (mean particle size 31 ± 30 μm, Meijer) was added to the solution, so that the final slurry was 70 vol % sucrose and 30 vol % polymer + nanoparticles. This was followed by addition of NaCl (Jade Scientific) at 60 vol% of the total polymer + nanoparticle volume to create macroporosity. The suspension was vortexed for 10 minutes and pressed into a silicon mold that was 4.7 mm diameter, 2 mm high. After air drying, devices were removed and washed for 2 hours in distilled water, changing the water every 30 minutes to remove sucrose and NaCl. After air drying devices overnight, they were stored in a desiccator prior to use. When nanoparticles were present, incorporation was confirmed via thermogravimetric analysis (Supplemental).

Porous films were formed from PCL and PLGA 50:50. Prior to casting films, polymers were solubilized in dichloromethane (DCM, Sigma); PCL was used at 4wt %, while PLGA solutions were at 10wt%. Again, sucrose was incorporated into the solution, so that the final slurry was 70 vol % sucrose and 30 vol % polymer. Suspensions were vortexed, then cast and dried on glass sheets. Dried films were removed and submerged in Milli-Q water to remove the sucrose, then dried for storage. In addition, non-porous PCL films were created with the same method but without the addition of nanoparticles or sucrose.

### 2.4 Cell studies

Primary human adult dermal fibroblasts were purchased from ATCC (cat no PCS-201-012), and used before passage 5 in all experiments. Fibroblasts were cultured in complete media (Dulbecco’s modified Eagle’s medium (DMEM, ThermoFisher Scientific, 11-965-092), 10% fetal bovine serum (FBS, Life Technologies, A5669501), and 1% Penicillin– Streptomycin (cat no 1510122)). As a positive control for myofibroblast differentiation, cells were cultured in differentiation media: complete media supplemented with 10 ng/ml tissue growth factor beta one (TGFβ1, Peprotech, cat 100-21-10UG).

#### Device culture

Dry devices were sterilized individually, by immersing the device in 70% ethanol for 60 min. In addition, PCL devices were centrifuged for 5 minutes at 11,000 rpm at the start of incubation. Following ethanol soaking, devices were washed in sterile phosphate buffered saline (PBS) and centrifuged for 10 minutes at 11,000 rpm. The devices were stored in fresh sterile PBS until use. An hour prior to use, devices were immersed in complete media.

For cell seeding, sterilized devices were placed in dry individual wells of a 48 well plate (non-treated). Fibroblasts (1×10^5^ cells) were pipetted onto the devices in 5μl media. The cells were placed in an incubator at 37°C, 5% CO_2_ for 4 hours. At that time, 400 μl of complete media was added to the well, submerging the device and cells. For the positive control, cells received 400 μl of differentiation media instead. Media was changed 3 times per week for the duration of the study. Samples were harvested at day 1, day 14 and day 28 for mechanical testing and cell number quantification. Protein expression was evaluated after 14 and 28 days.

#### Film culture

The films used for culture were composed of either PCL or PLGA 50:50 with 0wt% TaO_x_. Dry films were cut into 13mm diameter disks and were sterilized by immersing in 70% ethanol for 30 min, followed by two washes with sterile water. For in vitro cell tests, films were placed into 24-well inserts (CellCrown™, Z681903-12 EA). To reduce the leakage of media around the insert, each sample consisted of two layers: a bottom layer of non-porous PCL (cut to a circle of 24mm diameter) and a top layer of porous film (diameter: 13 mm). After assembly into the inserts, they were placed at the bottom of a 24 well tissue culture plate, and the entire assembly was sterilized for 30 min under UV light. Sterile inserts were stored at room temperature in a biological cabinet, in sterile water, until use.

Prior to the start of culture, sterile inserts were incubated with 200 μl of complete media for one hour. After removing the media, each well was seeded with 5 ×10^4^ fibroblast cells in 400 μl complete media. Cells were placed in an incubator (37°C, 5% CO_2_) for 4 hours to allow attachment. Then the media was changed to 400 μl complete media with 0, 0.01, 0.1, 1 or 10 mM lactic acid (Sigma Aldrich, cat no 252476). As a positive control, cells were treated with differentiation media. For all tests with lactic acid and the positive control, cells were cultured on PCL films; an additional group was cultured on PLGA 50:50 films with 0 mM lactic acid. After two weeks, films were harvested for quantification of protein expression.

### 2.5 Mechanical testing

The compressive modulus of devices with cells was measured at day 0, day 14 and day 28. In all cases, devices without cells (incubated under the same conditions as other samples) were used as controls to assess the effects of cellular activity. Prior to performing mechanical tests, device thickness and diameter was measured using a micrometer. Devices were placed in a 35mm Petri dish containing 2 ml PBS, situated within the mechanical tester (TAXT plus Texture analyzer, Stable Micro Systems, 5kg load cell). A glass coverslip was placed over the top of the sample before compressing to 40% of their original thickness, at 1 mm/min and then unloading. The start of the loading cycle was defined as when the force of 0.03N was reached. The loading cycle was repeated twice. The compressive modulus was calculated as the initial slope of the stress-strain curve from the first loading cycle. The percentage of deformation was obtained by dividing the device thickness after the initial loading cycle by the original thickness × 100%.

After testing, devices were removed from the machine, and placed in the microcentrifuge tubes on ice. After all samples were tested, they were washed in 1 ml deionized water for 5 minutes. The water was subsequently removed and the devices were frozen at −80°C, in preparation for cell proliferation testing. Results are presented as mean ± standard deviation.

### 2.6 Cell Attachment and Proliferation

To quantify attachment and proliferation, devices were harvested after 24 hours, and on days 14 and 28. Samples underwent mechanical testing and then were stored at −80 °C for DNA quantification using Quant-iT™ PicoGreen™ dsDNA Assay Kit (ThermoFisher Scientific), per manufacturer’s instructions. Frozen inserts were digested in 200 μl papain buffer (0.1M phosphate buffer, 10mM L-cysteine, 2mM Ethylenediaminetetraacetic acid (EDTA), 3U/ml papain) at 60 °C overnight. In the assay, samples were diluted 1:4 in TNE buffer and loaded on a black 96 well plate. An equal amount of PicoGreen dye (1:200 in TNE buffer) was added to each well, and fluorescence was read at Ex: 480 nm, Em: 520 nm using a BioTek plate reader. Two standards were used: a cell standard (0–5×10^4^ cells) of fibroblast cells, created on the day of experimental cell seeding, and a DNA standard (1 μg/ml - 1 mg/ml) provided by the kit. To ensure the reading was not caused by background fluorescence from the polymer inserts, devices incubated for the same amount of time in complete media but without cell culture were run and data was normalized to these blanks. All tests were done in triplicate, with two technical replicates each, and presented as mean ± standard deviation.

### 2.7 Protein Expression

Protein was harvested from devices and films in Ripa buffer. Protein concentration was measured (BioRad, BCA kit) and expression of myofibroblast markers was quantified using sodium dodecyl sulfate polyacrylamide gel electrophoresis (SDS-PAGE) and Western blotting. Full protocol details and images of complete blots are presented in Supplemental data. Briefly, samples with 2× Laemmli buffer were incubated at 60°C for 20 min, and separated by SDS/ PAGE. After transfer to PVDF membranes (iBlot 2 Gel Transfer Device, Invitrogen), membranes were blocked with 5% dried milk in PBST (phosphate buffered saline pH 7.4, containing 0.1% Tween-20) and probed with primary antibodies specific for alpha smooth muscle actin (αSMA, 1:1000, Abcam, ab7817), collagen Type I (1:1000, Life Technologies, PA595137), integrin α_V_ (1:1000, Abcam, ab302640), integrin β1 (1:1500, Abcam, ab179471), integrin β5 (1:500, Invitrogen, PA5-50991), and vinculin (1:10000, Abcam, ab129002) at 4 °C overnight. The next day, the membrane was washed and incubated with anti-mouse IgG-HRP (Millipore, 12-349) or anti-rabbit IgG-HRP (ab205718, Abcam) at room temperature for 2 hr. Protein bands were visualized with Amersham ECL Plus substrate (cytiva, RPN2232) on a Li-Cor Odyssey FC and resulting signal was quantified via Image J.

Developed membranes were washed in PBS and stripped for 15 minutes in Restore Western Blot Stripping Buffer (Thermo Scientific 21059) before being either blocked and reprobed, or stained for total protein using the BLOT-Fast Stain (G Biosciences, cat# 786-34) according to manufacturer’s directions. The total protein signal was imaged under visible light on a c300 imager (Azure biosystems). Protein expression was quantified via Image J and normalized to total protein to adjust the measured band intensity. For comparisons between membranes, all signals were normalized to PCL + 0wt% TaO_x_, which was run on every membrane. All data reported are the result of three independent replicates (two technical replicates each) and are reported as mean ± standard error.

### 2.8 Immunohistochemistry

Prior to staining, fixed samples were stored in PBS at 4 °C. Films were stained without further preparation. Devices were embedded in Tissue Tek Optimal Cutting Temperature (OCT) Compound and frozen on dry ice. Once embedded, a Leica CM3050S cryostat was used to make 20μm slices through the cross-section of devices and adhered to glass slides. Slides were allowed to air dry, and stored at −20°C until staining.

Prepared samples, either films or cryosectioned slices on slides, were washed in PBST, then permeabilized in 0.1% Triton X-100 in PBS, followed by two PBST washes. Samples were then blocked with 5 wt% bovine serum albumin (BSA) in PBST, for 1 hr, at room temperature. After washing twice in PBST, samples were incubated with primary antibodies, overnight at 4 °C, in 3 wt% BSA/PBST. Following primary incubation and two PBST washes, secondary antibodies in PBS were incubated with the samples for 1–2 h at room temperature. To visualize the actin cytoskeleton, samples were incubated for 30 minutes in ActinRed^TM^ 555 ReadyProbes^TM^ (Invitrogen, R37112, 2 drops per ml PBS). Finally, samples were washed twice with PBS and left in a solution of PBS and DAPI stain (1:000 in PBS, ThermoFisher Scientific, 62248). Primary antibodies, and the dilutions used, were: αSMA (1:300, Abcam, ab7817) and collagen Type I (1:500, Life Technologies, PA595137). Secondary antibodies: Alexa Fluor 488 (1:1000, A11070), and Alexa Fluor 647 (1:1000. A21236).

Stained samples were inverted onto a coverslip in a drop of Slow Fade Diamond Antifade Mountant (Invitrogen, S36972). Imaging was performed using a Leica DMi8 microscope, using an LASX software interface. With the high surface roughness of the samples, z-stacks were taken at a minimum of 3 places and post-processed with Thunder image analysis (Leica) to remove background fluorescence.

### 2.9 Device imaging

Micro-computed tomography (µCT) imaging was performed on a Revvity Quantum GX. Devices were imaged in PBS at 90 keV, 88 µA, with a 25 mm field of view at a 50 µm resolution. Samples were imaged while immersed in cell culture media, in individual wells of a 48 well plate.

Device architecture was imaged via scanning electron microscopy (SEM). Cryosectioned slides were immersed in PBS at room temperature for 5 minutes, in 70% ethanol for 5 minutes and then washed in 100% ethanol and allowed to air dry to remove OCT compound. After drying, sections were transferred to 13mm aluminum stubs with double sided carbon tape. Stubs were then sputter coated with platinum and examined using a MIRA3 Tescan, at 10 keV in scanning electron mode.

### 2.10 Statistics

Statistics were performed using GraphPad 10.4.0. Groups were compared via ANOVA, followed by Fisher’s LSD test. In all cases, α < 0.05 was considered significant, with a 95% confidence interval.

## 3. Results & Discussion

Foreign body reactions to biomaterials are complex and often determine whether implanted devices are clinically successful or not [11]. In some of the worst cases, the initial inflammatory reaction post-implantation leads to peri-implant fibrosis, or fibrotic encapsulation, with a dense layer of extra-cellular matrix (ECM). When this occurs, devices cannot fulfill their primary function and must be removed, increasing patient morbidity [2].

Both macrophages and fibroblasts are essential in the foreign body response, often working in tandem [12]. The fibroblast transition to a myofibroblast phenotype presages encapsulation, although there are still many open questions surrounding the origin and sub-populations of fibroblasts that respond in foreign body reactions [1,4]. Given their role in peri-implant fibrosis, it is important to understand how fibroblasts react to material properties within tissue engineering devices. This class of biomedical devices are microporous and often made from polymers, with properties that are tuned via matrix chemistry or architectural features. In this study we concentrated on the effects of matrix chemistry, focusing on nanoparticle additions to a polymer matrix and the effect of the matrix itself for driving a transition to a myofibroblast phenotype.

Nanoparticle additions to polymer matrices have been employed for a variety of purposes, such as carriers for therapeutic delivery [13], as antimicrobial agents [14], or as contrast agents for medical imaging [15]. In all cases, nanoparticles must be biocompatible, preferably eliciting a minimal inflammatory response when used in implanted devices, and have the ability to be homogeneously incorporated within the polymer. Therefore, a biocompatible tantalum oxide (TaO_x_) nanoparticle was chosen. It has been demonstrated to confer radiopacity when incorporated into radiolucent polymers [8,9], and at up to 20wt% tantalum incorporation has shown only minor inflammatory responses from macrophages [10]. While a polyester matrix was used in all cases, the polymers chosen (PCL and PLGA) exhibit a range of mechanical properties and degradation profiles. This allowed for a more in-depth investigation into the effect of each variable on myofibroblast differentiation. Regardless of the polyester, or nanoparticle addition, the porous architecture of the tissue engineering device mimics was kept constant, as the structure of devices also affects the foreign body reaction independent of material [12]. Devices, seen in Figure 1(a), consisted of a macroporous architecture with mean pore size of 350-400 μm (Fig 1(b)), and microporous walls (Fig 1(c)) with thin struts between pores (Fig 1(d)). The range of features accommodates both nutrient diffusion and cellular attachment, and fibroblasts were able to proliferate within the structure over four weeks of culture, Figure 1(e-f).

**Figure 1:**
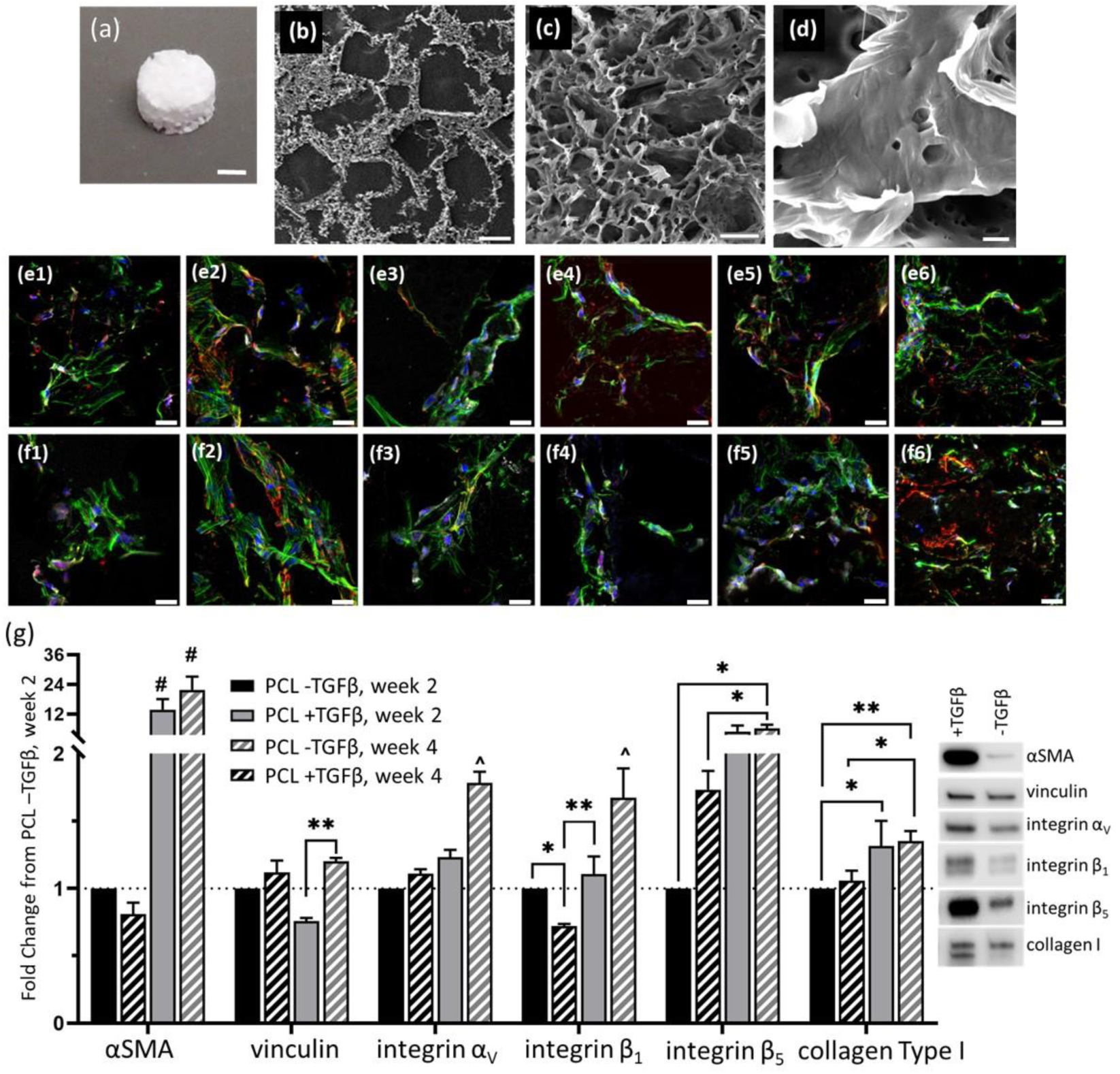
Porous tissue engineering devices supported growth and differentiation of human adult dermal fibroblasts. (a) Seen macroscopically, devices were designed to mimic tissue engineering implants. With electron microscopy, hierarchical features were visible, including (b) macroporosity, (c) microporosity in the device walls and (d) individual struts of the micropores. The features allowed fibroblasts to grow into the devices at (e) 2 weeks and (f) four weeks; red: αSMA, green: actin, white: collagen Type I, blue: nucleus. 1: PCL, 2: PCL + TGFβ1, 3: PCL + 10wt% TaO_x_ nanoparticles, 4: PCL + 20wt% TaO_x_ nanoparticles, 5: PLGA 85:15, 6: PLGA 50:50. (g) Fibroblasts were capable of transitioning to myofibroblasts, common to fibrotic phenotypes, as shown by protein expression over four weeks, quantified by Western blotting. All expression data was normalized to PCL -TGFβ at week 2, and protein loading on the blot was normalized by total protein loaded (supplemental). Protein expression data is presented as mean ± standard error and characteristic bands at week 4 are shown. α < 0.05, * p < 0.05, ** p < 0.01, # significantly different from all other groups (p < 0.0001), ^ significantly different from all other groups (p < 0.001). Scale bar (a) 1 mm, (b) 250 μm (c) 15 μm, (d) 1 μm, (e-f) 25 μm.

Importantly, it was confirmed that in vitro culture on porous devices supported fibroblast differentiation towards a myofibroblast phenotype, a hallmark of fibrotic diseases, with the addition of 10 ng/ml TGFβ1 in the culture media, Figure 1(g). As expected, this led to a significant increase in αSMA starting at week two and continuing throughout the culture period. In addition, by week four, supplementation led to a significant increase in collagen Type I expression, and expression of attachment integrins α_V_, β_1_and β_5_.

In fibrotic tissue, TGFβ1 is activated by integrins: α_V_β_6_ in lung and α_V_β_5_ in skin [16,17]. It has been hypothesized that fibroblasts express the integrin α_V_ sub-unit in excess and control myofibroblast differentiation through expression of the β sub-unit [17]. In the present study, stimulation of myofibroblast differentiation correlated to an up-regulation of both the β_1_and β_5_ sub-unit, which may reflect the cellular machinery required to attach to the polyester substrate. The effect on vinculin expression, present in focal adhesions and related to mechanical force generation in fibroblasts, was less pronounced [18].

### 3.1 Cell activity affects device properties

The foreign body reaction is dependent on the tissue microenvironment. In general, increased fibroblast attachment in vitro equates to greater capsule formation around devices in vivo [19]. There are a number of cues inherent in biomedical devices that can affect fibroblast attachment and differentiation, including mechanics, topography, surface chemistry and bulk material chemistry [4].

In the current study, there were no significant differences in the initial attachment of human adult dermal fibroblasts due to polymer matrix or nanoparticle content 24 hours after seeding. However, the ability for fibroblasts to proliferate within the porous structure was significantly impacted by matrix properties. The greatest proliferation was observed on matrices that were stable over time, namely those based on the non-degrading PCL [20]. In addition, over 4 weeks in culture, there was a significant increase in cell proliferation with increasing TaO_x_ nanoparticle content, Figure 2(a). Devices of PCL with 20wt% TaO_x_ had an average of 611.1 ± 82% growth, compared to PCL without nanoparticles at 208.6 ± 109% growth. On PLGA devices, a polyester with increased degradation compared to PCL, cell proliferation occurred over the first two weeks of culture and then cell number remained constant. Stimulating the myofibroblast transition with the addition of TGFβ1 appeared to promote greater cell proliferation over time, but the trend was not significant. While cells could penetrate through the devices, most remained near the top and side surfaces (supplemental).

**Figure 2:**
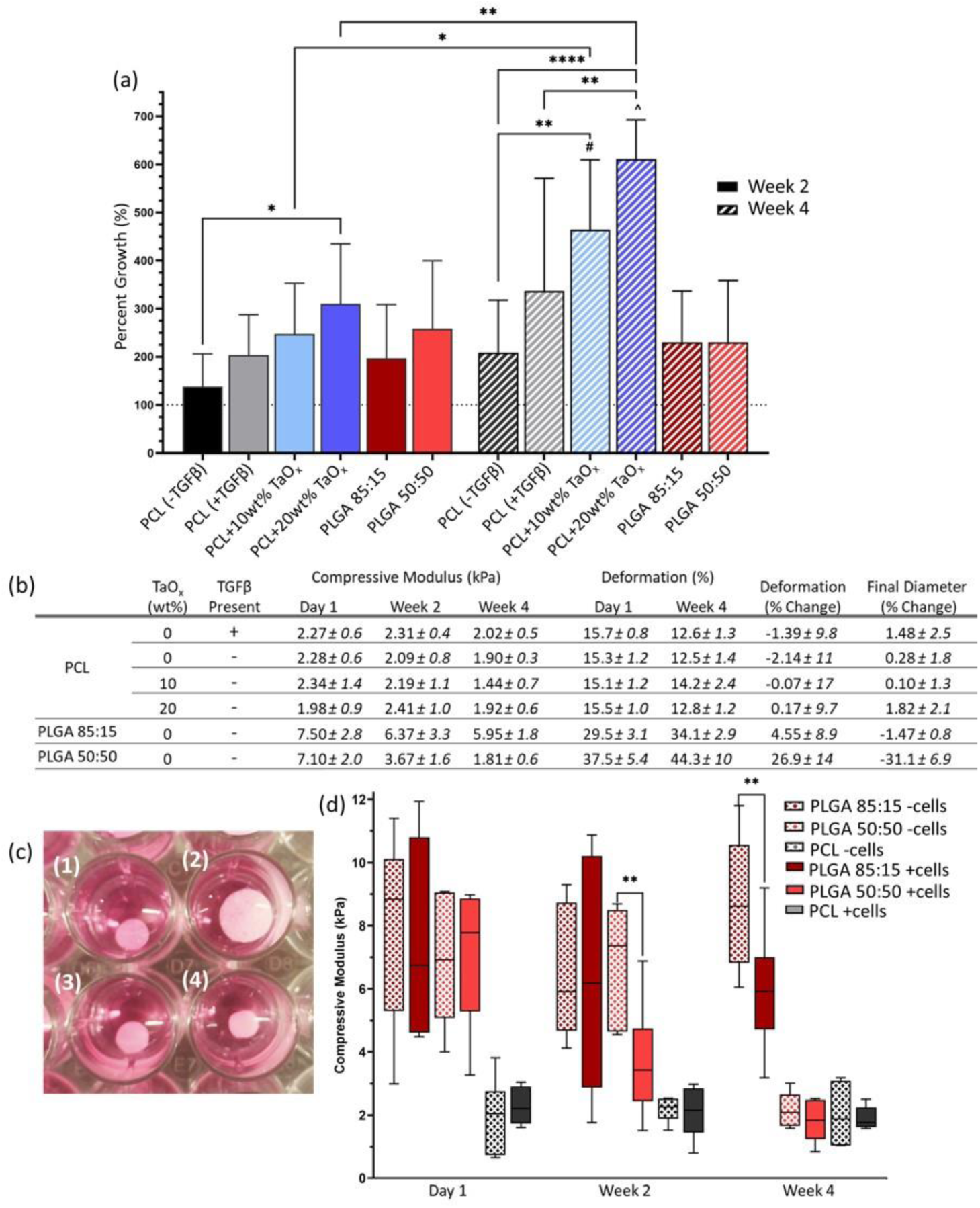
Device proliferation and mechanical properties are determined by the device matrix and can be affected by cellular interactions. (a) Percent growth over 4 weeks was significantly higher on composites of PCL + nanoparticles than PLGA matrices (mean ± st deviation). (b) Table of mechanical properties measured over four weeks. Compressive modulus and permanent deformation were measured on devices in contact with fibroblasts; the percentage change in deformation and final diameter are reported as a percent of control devices without cell culture. (c) Macroscopic effects on the device dimensions were noted due to cell interactions: (1) PLGA 85:15 without fibroblasts, (2) PLGA 50:50 without fibroblasts, (3) PLGA 85:15 with fibroblasts, (4) PLGA 50:50 with fibroblasts. (d) Cell interactions led to significantly faster degradation, measured via the compressive modulus, for PLGA devices; box plot whiskers show the spread of data from minimum to maximum values. α < 0.05, * p < 0.05, ** p < 0.01, ****p < 0.0001

Culturing fibroblasts within the devices significantly affected the device properties over two weeks. This is in agreement with literature that highlights increased degradation of polymer devices implanted in vivo compared to acellular in vitro environments, hypothesized to be due to increased fluid flows and protease concentrations [10,21]. Macroscopic changes to device dimensions were observed over four weeks of culture, Figure 2(b-c). This was particularly true for PLGA 50:50 devices, that have the fastest degradation time of the polyesters studied, degrading within a matter of weeks [9]. While the percent change from the original diameter was mild for PLGA devices (−6.10 ± 9.4%), since swelling was observed in the PLGA 50:50 control without cells, it correlated to a −31.1±6.9% change in final diameter between devices with and without cells, Figure 2(b).

In addition to observing dimensional changes, matrix mechanics, including compression and deformation, were monitored over in vitro culture. Overall, PCL device mechanics were not significantly affected by cell culture over time for any condition tested. Even TGFβ1 supplementation that drove myofibroblast transition did not affect macroscale mechanics. On the other hand, PLGA devices were significantly affected by fibroblast culture, Figure 2(a,c). Namely, devices lost significant structural integrity and mechanical properties up to two weeks earlier when cells were present, compared to controls that were incubated in media without cells, Figure 2(d). By the end of four weeks, the compressive modulus of PLGA 50:50 was not significantly different than PCL, 1.81 ± 0.6 kPa and 1.90 ± 0.3 kPa, respectively. PLGA 85:15, with its slower degradation rate than PLGA 50:50, had only begun to lose structural integrity at four weeks of in vitro culture. However, the permanent deformation in PLGA devices increased over time regardless of polymer blend. At four weeks, deformation reached 34.1 ± 2.9% and 44.3 ± 10% for PLGA 85:15 and PLGA 50:50, significantly greater than the initial deformation of devices observed after 24 hours.

PLGA is known to be sensitive to the action of proteases, which are present in complete culture media, and their expression is triggered during the foreign body response [22]. With the presence of fibroblasts, polymer chain scission was increased, observed as a significant decrease in mechanical strength of devices with cells compared to acellular counterparts. In addition, swelling of the PLGA devices was decreased by 31% with fibroblasts present in the matrix. Swelling or contraction of PLGA devices during degradation is significantly affected by pH conditions [9]. Matrix morphology can also be altered by cells exerting contractile forces on the matrix over time. This phenomenon has been most visible in devices with low mechanical strength, such as collagen implants [23]. However, the role of myofibroblast activity on contraction is believed to be relatively minor, as the induction of myofibroblast differentiation on PCL was not enough to alter the macroscopic device mechanics over the four weeks of culture.

### 3.2 Nanoparticle incorporation has a limited effect on fibroblasts

Composite materials based on polymers, have a long history of use as biomedical devices. Nanoparticles are increasingly being incorporated into polymers to provide added functionality. In the present case, TaO_x_ nanoparticles introduced into a PCL matrix conferred radiopacity to the device, Figure 3(a), that scaled with nanoparticle content, as noted in literature [10,24]. While fibroblast proliferation increased with TaO_x_ nanoparticles, they had very little effect on the protein expression of fibroblasts. At 20wt%, nanoparticles significantly reduced the expression of key myofibroblast markers over four weeks, Figure 3(b-d). In particular, the expression of αSMA was significantly down-regulated, reaching 0.31 ± 0.1 fold change from PCL + 0wt% TaO_x_ at week two. Integrin α_V_ expression was also sensitive to nanoparticle addition, leading to significant down-regulation. No significant changes were noted in vinculin, integrin β_1_ or collagen Type I expression.

**Figure 3:**
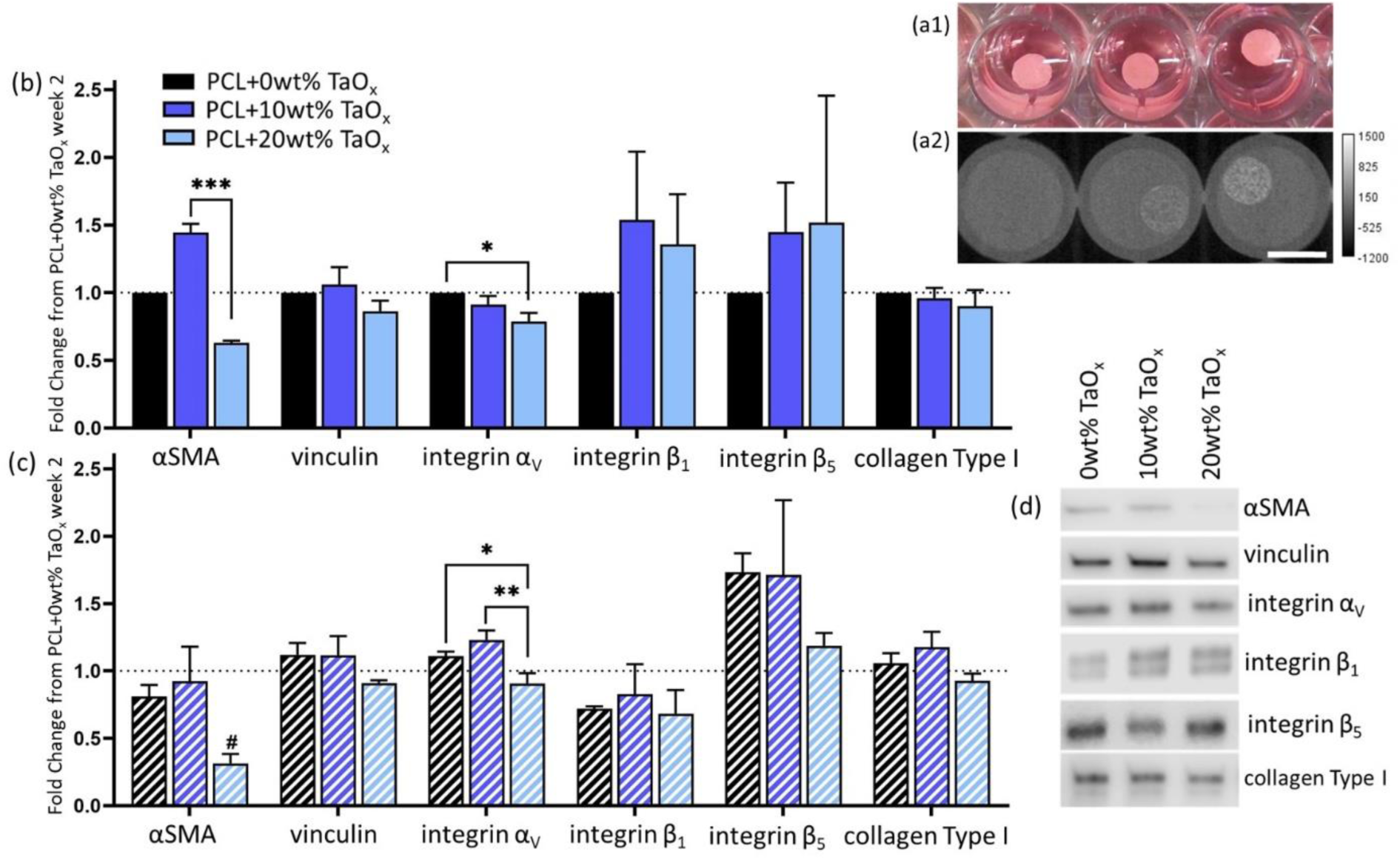
Nanoparticle addition in PCL devices did not promote myofibroblast differentiation. (a) After four weeks in culture, PCL+ radiopaque TaO_x_ devices remained intact, seen (1) visually and (2) via μCT. TaO_x_ nanoparticle addition increased radiopacity of devices; from right to left: PCL + 0wt% TaO_x_, PCL + 10wt% TaO_x_, PCL + 20wt% TaO_x_. Protein expression of myofibroblast markers were either stable or down regulated with increasing TaO_x_ addition after (b) two weeks or (c) four weeks. (d) Typical Western blotting protein bands at week four for markers of myofibroblast transition: αSMA, vinculin, integrin α_V_, integrin β_1_, integrin β_5_, and collagen Type I. All expression data was normalized to PCL + 0wt% TaO_x_ (without TGFβ1 supplementation) at week 2, and protein loading on the membrane was normalized by total protein (supplemental). Protein expression data is presented as mean ± standard error. α < 0.05, * p < 0.05, ** p < 0.01, # significantly different from all other groups at this time point (p < 0.05). Scale bar: (a) 5 mm.

In other cell types, the biological response associated with the introduction of TaO_x_ nanoparticles has been mediated by changes to the surface topography of the composite, namely increased nanoscale surface roughness [10,25]. In literature, nanoscale surface roughness has been linked to decreased fibrous capsule formation and collagen deposition compared to devices with predominately microscale features [26,27]. The reduced fibrotic deposition around implanted devices has been linked, not only to increased device survival, but to increased density of blood vessels [2,27].

It is not clear if the decreased expression of myofibroblast markers is linked with altered cell attachment mechanisms. In studies on silicone implants, blocking integrin α_V_ reduced fibroblast activation by reducing sensitivity to substrate mechanics [28]. However, the effects of surface topography may also be rooted in changes in wettability, as more hydrophilic surfaces drive the myofibroblast transition [29]. In all cases, altered foreign body responses are likely mediated by changes in protein adsorption with surface topography, with lower adsorption leading to a limited response [4].

### 3.3 Degradation products drive cellular behavior

Polyester matrices are very versatile, representing a wide range of mechanical properties and degradation profiles, but also a wide range of foreign body responses. In concurrence with their known biocompatibility, fibroblasts attached and proliferated on all matrices tested, Figure 4(a). While nanoparticle addition had a limited effect myofibroblast markers, properties of the polymer matrix stimulated greater changes in protein expression. Of note, αSMA expression was significantly increased over four weeks on PLGA 50:50 devices, reaching a fold change of 5.16 ± 2.5 compared to PCL. Cells on PLGA 85:15 showed a 2.5 ± 1.5-fold up-regulation, Figure 4(b-d). In addition, PLGA 50:50 also stimulated higher expression of integrin β_1_and integrin β_5_. However, at no point was marker expression of αSMA, or integrin β_5_on PLGA 50:50 higher than the positive myofibroblast differentiation control (PCL + TGFβ). These results track with literature where PCL and PLGA matrices implanted in vivo have been shown to trigger different immune responses [21].

**Figure 4:**
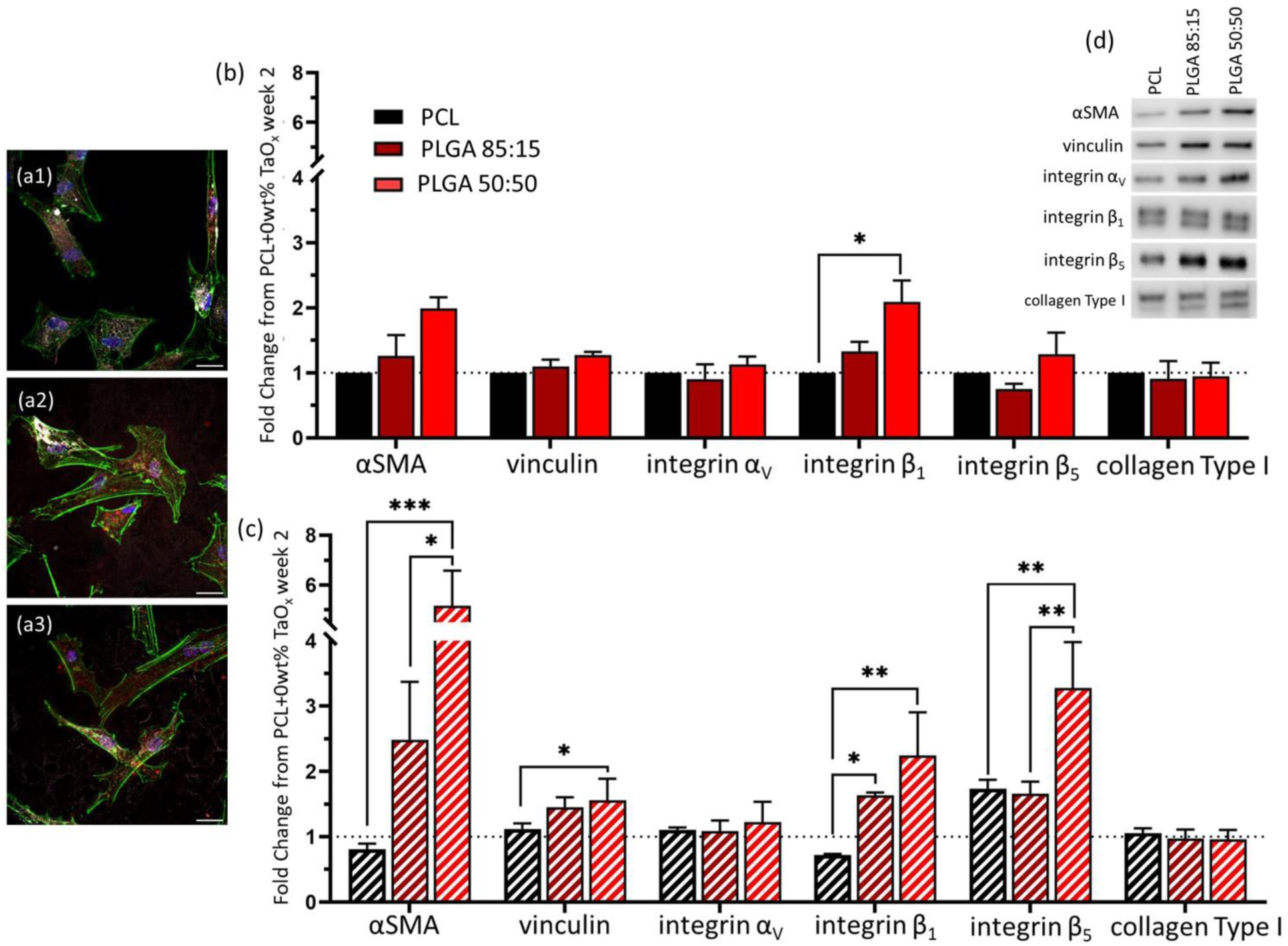
The polymer matrix making up devices had a significant impact on myofibroblast marker expression over four weeks. (a) Fibroblasts were attached and well spread on the surface of all polymers after one week culture on films (1) PCL, (2) PLGA 85:15 (3) PLGA 50:50. Green: actin, red: αSMA, white: collagen Type I, blue: nucleus (DAPI). Protein expression was quantified via Western blotting at (b) two weeks and (c) four weeks of culture. Markers examined were αSMA, vinculin, integrin α_V_, integrin β_1_, integrin β_5_, and collagen Type I. All expression data was normalized to PCL (+ 0wt% TaO_x_, without TGFβ1 supplementation) at week 2, and protein loading on the membrane was normalized by total protein (supplemental). (d) Typical protein bands of markers at four weeks. Protein expression data is presented as mean ± standard error. α < 0.05, * p < 0.05, ** p < 0.01, *** p < 0.001. Scale bar: (a) 25 μm.

Fibroblasts are able to respond to mechanical cues within their environment and can respond to changes over several orders of magnitude [28]. The range of apparent compressive modulus of the devices used in the present study were between 2-7 kPa, Figure 2(b). Even in this range, in line with the modulus of natural tissue, greater mechanical properties increase αSMA expression and vinculin organization [30]. It should be noted that the apparent modulus reflects a combination of the inherent compressive modulus of the polymer and characteristics of the device architecture. Thus, fibroblasts cultured within the devices may be perceiving much greater mechanical forces than recorded during testing.

However, mechanical properties of the matrix cannot explain the observed changes in protein expression. Both PLGA 85:15 and PLGA 50:50 began the study with comparable compressive moduli, and yet αSMA expression on PLGA 50:50 devices only significantly increased after mechanical strength was lost. As noted in Figure 2, by four weeks, the mechanical strength of PLGA 50:50 had fallen to the same level as PCL, and yet marker expression was significantly greater. It is more likely that the release of degradation products, such as lactic acid, drive protein expression, as mass loss begins after mechanical modulus decreases [31]. Of the polyesters studied, PLGA 50:50 has the fastest degradation rate, followed by PLGA 85:15 [9]. It is noted that marker expression on PLGA 85:15 appeared to be following the same trends as PLGA 50:50, but at lower rates, consistent with a lower level of degradation products.

Degradation products, and lactic acid in particular, is believed to mediate the biological response to polymers in many systems, including glia and macrophages [32,33]. Lactic acid is part of the glycolytic pathway and can shift cellular metabolism in ways that up-regulate pro-inflammatory factors [33]. Glycolysis metabolism may also lead to an up-regulation of reactive oxygen species that induce TGFβ1 gene expression in fibroblasts, ultimately inducing fibrosis [34]. To further test the hypothesis that lactic acid was responsible for the protein expression changes, fibroblasts were cultured on PCL microporous films, the substrate with the lowest potential for inducing myofibroblast differentiation, with lactic acid supplementation in the media. The films were designed to mimic the microporous device walls, without creating gradients of oxygen or metabolites induced within 3D devices, to ensure all cells experienced elevated lactic acid levels.

With the addition of lactic acid to the culture media, the expression of fibroblast markers changed significantly, Figure 5(a-b), but cells remained well spread and attached to the PCL substrates (Fig 5(c)). Moderate doses of 0.1 - 1 mM lactic acid had the greatest positive effect on the expression of all proteins at week 2. At 10 mM, the expression of myofibroblast markers was reduced back to baseline levels, Figure 5(a-b). Films of PLGA 50:50 have been shown to have a burst release of lactic acid and di-lactic acid over the first week, reaching a cumulative release of 0.015 mM in the absence of proteases [32]. It is likely that elevated protease activity during in vitro cell culture increases the release rate of lactic acid, potentially bringing the concentration of lactic acid within the range that stimulated the greatest myofibroblast activation.

**Figure 5:**
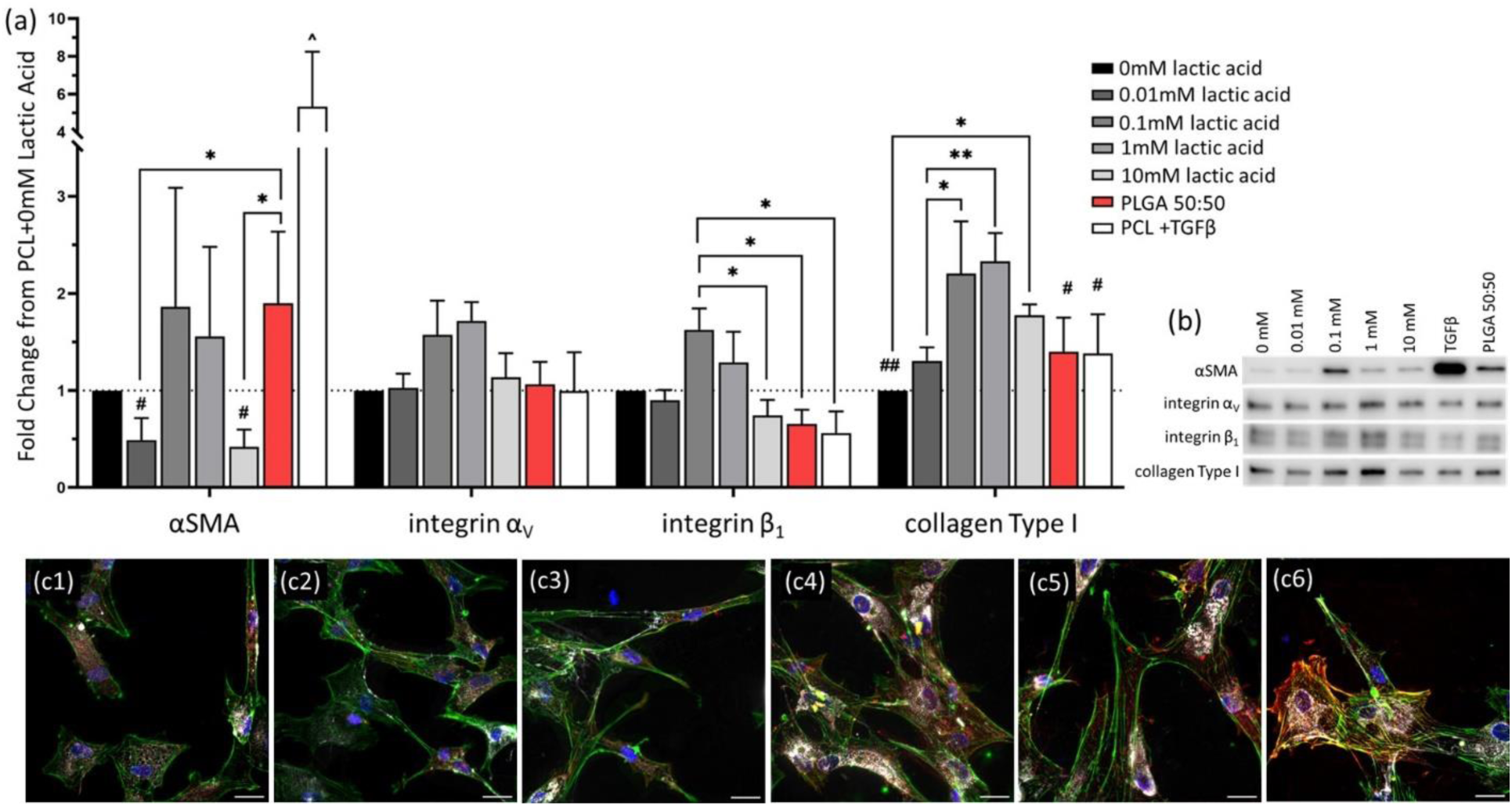
Degradation products of PLGA matrices, namely lactic acid, significantly impact fibroblasts. (a) Protein expression of fibroblasts, on microporous films after two weeks of culture, was altered by addition of 0 - 10 mM lactic acid, as seen in (b) typical protein bands in Western blotting. All expression data was normalized to PCL without TGFβ1 supplementation (0 mM lactic acid), and protein loading on the membrane was normalized by total protein (supplemental). Protein expression data is presented as mean ± standard error. (c) Fibroblasts attached and were well spread regardless of media supplementation (1) 0 mM lactic acid, (2) 0.01 mM lactic acid, (3) 0.1 mM lactic acid, (4) 1 mM lactic acid, (5) 10 mM lactic acid, (6) 0mM lactic acid + TGFβ1 supplementation. Fibroblasts were stained for collagen Type I (white), αSMA (red), actin (green) and nucleus (blue). α < 0.05, * p < 0.05, ** p < 0.01, # significantly lower than 0.1-1 mM lactic acid (p < 0.05), ## significantly lower than 0.1-1 mM lactic acid (p < 0.01), ^ significantly greater than all other groups (p < 0.001). Scale bar: (c) 25 μm.

Once again, PLGA 50:50 films had increased αSMA expression compared to PCL films with no lactic acid supplementation. However, changes in attachment proteins on the films did not follow trends seen in three-dimensional devices. The transition from 2D to 3D is known to affect cell attachment mechanisms, making it likely that differences in expression were due to additional architectural cues or local pH conditions within the confines of the device that were not present during film culture [35].

Lactic acid has previously been shown to drive the myofibroblast transition in lung fibroblasts and in adipocytes, driven by changes in pH [36,37]. In these studies, 10 mM lactic acid supplementation gave the same marker expression as TGFβ1 after three days of culture [36]. It is hypothesized that this effect might be transitory and have peaked by the two week time point used in the present study, which was required to get comparable degradation of PLGA 50:50. The use of a microporous film substrate, compared to a traditional tissue culture polystyrene or glass may also affect protein expression of cells in culture, by altering cell shape and attachment machinery.

In addition to in vitro effects, lactic acid has been associated with pathological conditions related to fibrosis [36]. In clinical patients, elevated levels of lactate have been found in lung tissue of patients with idiopathic pulmonary fibrosis (IPF) and in the blister fluid of scleroderma patients [34,38]. This suggests that the preclinical and in vitro results regarding the effects of lactic acid can be translated to humans. Indeed, PLGA 50:50 is known to stimulate a robust inflammatory response after implantation, although no long-term complications have been reported for clinical devices [22,39]

Ideally, biomedical implants can be engineered to encourage the attachment of fibroblasts to anchor implants, while avoiding the processes that lead to encapsulation [29]. The ability for degradation products to stimulate biological responses is well recognized [40]. However, it is important to realize that material degradation products can affect the foreign body reaction even at the initial moments of implantation. A limitation of this study is its focus on in vitro responses. Within the in vivo environment, micro-mechanical forces play a role in determining the severity of response to implanted devices across species [41], and it is becoming increasingly apparent that the biological milieu around the device can also affect the type and extent of the foreign body reaction [24,42]. Despite the simplified system, this study serves as a starting point for discussion around the contribution of material factors within the foreign body reaction.

With peri-implant fibrosis playing a large role in device failure, strategies have been proposed to mitigate this, including the incorporation of anti-inflammatory therapeutics into polymer matrices, such as capsaicin [2, 43]. Since myofibroblast differentiation requires lactate transporter activity and lactate dehydrogenase, inhibition of these proteins might also be good targets for a therapeutic strategy and are being tested for treating fibrotic disease [36, 44]. Whether tracking devices via in situ imaging, or delivering therapeutics to ensure tissue integration, nanoparticle incorporation into polymer matrices is poised to play a key role in the next generation of medical implants. From the current study, the use of nanoparticles may also have previously unappreciated effects and lead to a reduction in peri-implant fibrosis. Together these highlight the interplay of different device characteristics in determining the foreign body response.

## 4. Conclusions

Biomedical devices interact with native tissues via the foreign body response. While most literature has focused on the immune component of this process, this study emphasizes the role of fibroblasts in responding to biomaterials cues, such as polymer chemistry and surface topography. When human dermal fibroblasts were cultured on porous polymer devices, the presence of cells increased the degradation rate of polymer devices. The introduction of up to 20wt% TaO_x_ nanoparticles into the polymer matrix, rendered the device radiopaque, and led to a decrease in myofibroblast markers. The choice of polymer matrix had the most significant effect on myofibroblast activation. The fastest degrading polymer, PLGA 50:50, released lactic acid into the culture environment that could significantly up-regulate the protein markers for myofibroblasts, namely αSMA, reaching a 5.16 ± 2.5-fold increase over inert PCL matrices. The lactic acid released during degradation was shown to contribute to the expression of myofibroblast markers, separate from substrate mechanics. Together the addition of nanoparticles and tailored degradation of polymers are two ways that engineers can tune biomedical devices to avoid the excessive per-implant fibrosis that leads to clinical device failure.

## 5. Acknowledgments

The authors would like to thank Per Askeland from the MSU Composite Materials and Structures Center for help with scanning electron microscopy. This study was supported by the National Institute of Biomedical Imaging and Bioengineering of the NIH under award number R01EB029418. The content is solely the responsibility of the authors and does not necessarily represent the official views of the National Institutes of Health.

## Statements and Declarations

### Data availability

Data for this article, including data for cell proliferation, mechanics and protein expression are available at Mendeley Data at https://doi.org/10.17632/ynwydgkdz2.1

