## Supplemental for "Distinct Roles of Surface Nanostructure and Polymer Degradation in Fibroblast Response to Device Design"

### S1. $TaO_x$ Nanoparticle Characterization

Hydrophobic $TaO_x$ nanoparticles were manufactured based on our previous procedure with one minor change.[1] Hydrophobicity was imparted by coating nanoparticles with hexadecyltriethoxysilane (HDTES, Gelest Inc, cat no SIH5922.0). The $TaO_x$ nanoparticles were imaged via transmission electron microscopy, **Figure S1**. They were spherical particles, with a mean diameter of 6.5 ± 3nm when analyzed via Image J.

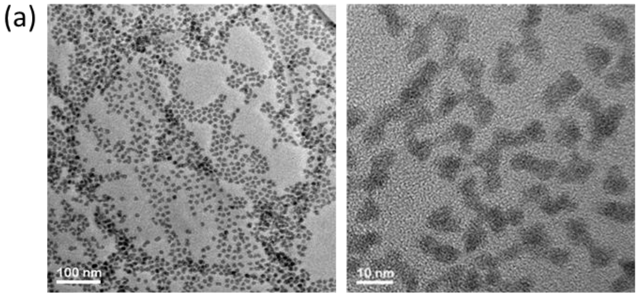


(b)

| Nominal $TaO_x$ (wt%) | Measured $TaO_x$ (wt%) |
|---|---|
| 0 | 0 |
| 10 | 8.878 |
| 20 | 16.138 |

**Figure S1**. Hydrophobic $TaO_x$ nanoparticle characterization. (a) Visualized via transmission electron microscopy, the particles show a tight distribution in size (3-9nm) at all magnifications. (b) The amount of nanoparticles incorporated within devices, measured via TGA, was between 10-15% of the nominal value.

Thermal gravimetric analysis (TGA) was used to characterize the wt% of $TaO_x$ nanoparticles in dichloromethane (DCM) after manufacture and the final wt% of $TaO_x$ incorporated into polymeric devices. To determine initial wt% $TaO_x$, the DCM solution was massed in an alumina pan and allowed to dry before placing in a TA Q500 (TA Instruments). For wt% $TaO_x$ in devices, 12-14 mg of dry material was massed in an alumina pan. Samples were then heated to 600°C at a rate of 10°C/min under a nitrogen environment. The percentage of nanoparticles was calculated as the difference between the initial and final mass; for solutions the initial mass was the mass of the DCM solution added to the pan.

### S2. Fibroblast Distribution within Devices

Devices cultured with fibroblasts for two and four weeks were fixed in 3.7% paraformaldehyde for 15 minutes. The devices were then incubated in phosphate buffered saline with 0.1% Tween-20 (PBST) for 30 minutes and permeabilized in 0.1wt% TritonX-100 for 10 minutes, while rotating. After 2 PBST washes, 5 minutes each, devices were blocked in 5wt% bovine serum albumin (BSA)/PBS for 2 hours. After 2 washes in phosphate buffered saline (PBS), the devices were incubated in HCS Nuclear Mask Red (1:1000 dilution, cat no 2738406) for 3 hours to stain cell nuclei. Devices were washed in PBS and frozen on dry ice in Tissue-Tek Optimal Cutting Temperature (OCT) Compound. Once frozen, the blocks were placed on an Emit Xerra. Images were taken of the block every 20µm. The slices were reconstructed into a three-dimensional view of the block with embedded devices. Representative cross-sections of individual devices were isolated using Image J, **Figure S2**.

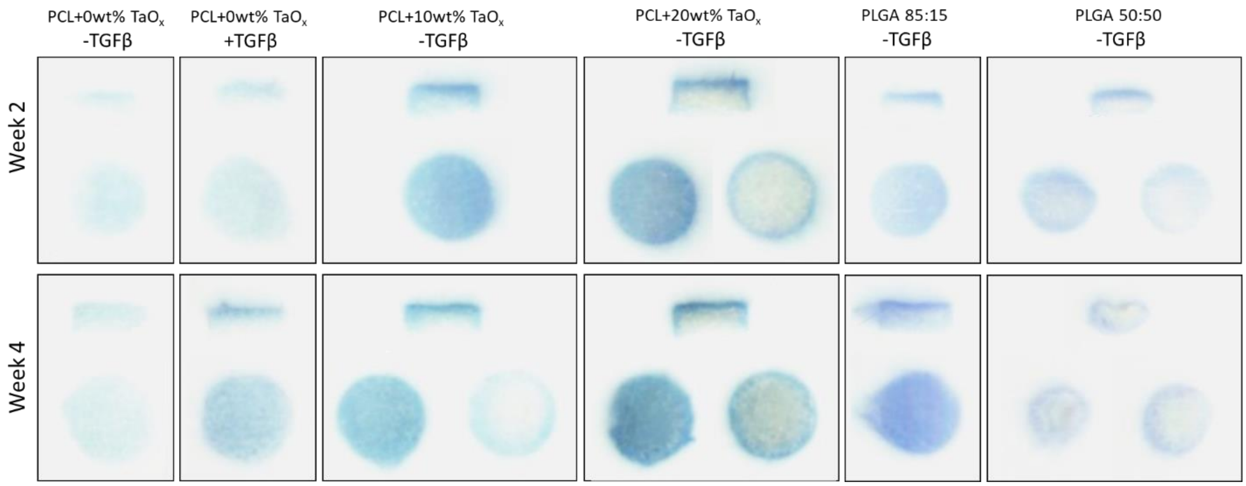


**Figure S2.** Human primary dermal fibroblasts were cultured on porous devices for up to four weeks and showed a typical pattern of proliferation, with the greatest number of cells at the top and sides of the devices. Cell nuclei were stained blue prior to sectioning. In all panels, the top image is a cross-section through the device thickness, with one or two slices parallel to the device top surface. All devices were originally the same size: 4.7mm diameter, 1.5mm thick.

**S3. Protein Expression**

Western blotting was used to quantify protein expression. Co-cultures were lysed in Ripa buffer (supplemented with phosphatase inhibitors and protease inhibitors), and spun down at 15,000×g for 5 min at 4 °C. After which, supernatants were collected and the protein concentration of the supernatant was measured (BioRad, BCA kit). Supernatants were combined with 2× Laemmli buffer, incubated at 60°F for 20 min, separated by SDS/ PAGE (4 μg of protein per lane), and transferred to PVDF membranes. Transfer was accomplished using the iBlot 2 Gel Transfer Device (Invitrogen, IB21001), a dry transfer system, at the recommended setting of 20 V for 1 min, 23 V for 4 min and 25 V for 2 minutes (7 min total). PVDF membranes were blocked with 5% dried milk in PBST (phosphate buffered saline pH 7.4, containing 0.1% Tween-20) and probed with primary antibodies specific for alpha smooth muscle actin (αSMA, 1:1,000, Abcam, ab7817), collagen Type I (1:1000; Life Technologies, PA595137), integrin $\alpha_V$ (1:1000, Abcam, ab302640), integrin $\beta_5$ (1:500, Invitrogen, PA5-50991), integrin β1 (1:1500, Abcam, ab179471), and vinculin (1:10000, Abcam, ab129002) at 4 °C overnight. The next day, the membrane was washed and incubated with anti-chicken IgY-HRP (12-341, Millipore) or anti-rabbit IgG-HRP (ab205718, abcam) at room temperature for 2 hr and further washed in PBST six times.

Protein bands were visualized with ECL Plus substrate (cytiva, RPN2232) on a Li-Cor Odyssey FC and resulting signal was quantified via Image J. Membranes were then washed in PBS and stripped for 15 minutes in Restore Western Blot Stripping Buffer (Thermo Scientific 21059) with subsequent PBS washing. Membranes were either blocked and reprobed or were stained for total protein using the BLOT-Fast Stain (G Biosciences, cat# 786-34) according to manufacturer's directions. The total protein signal was imaged under visible light on a c300 imager (azure biosystems).

Signal for total protein in each lane was quantified on Image J, and the protein loading was normalized to the signal from the first lane. The normalized signal was used to adjust the measured band intensity from antibody staining, also quantified using Image J. For comparison of nanoparticle effect and matrix protein, all protein expression was recorded as a fold change from PCL matrices, with no TGFβ supplementation, with 0wt% $TaO_x$ , at week 2. The final protein quantification reported is the result of three biological replicates (2 technical replicates each). Samples of PCL + 0wt% $TaO_x$ at week 2 were present on all blots and used to normalize differences between membrane exposure.

In the following figures (**Figure S3-S13**), each set of membranes is presented with all protein bands shown, along with the total protein staining for each membrane. For integrin and collagen Type I expression, only bands at high molecular weight, corresponding to the mature protein, were quantified. In cases where the protein ladder was disproportionately dark compared to the band signal, the molecular weight of bands was determined from an initial blot, with a molecular weight marker present, and the ladder was omitted in subsequent blots.

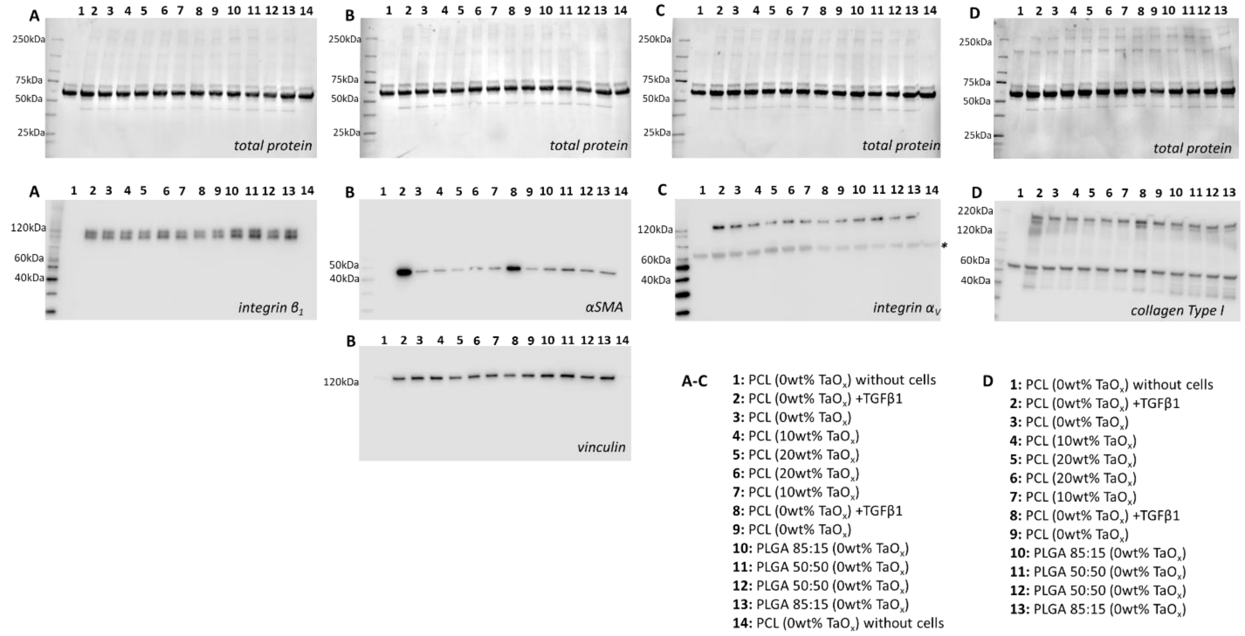


**Figure S3:** Western blots of porous devices at week 2 (biological replicate 1). Top row: total protein, second and third rows (A) integrin $\beta_1$, (B) $\alpha$SMA, vinculin, (C) integrin $\alpha_V$, (D) collagen Type I. The sample loaded in each lane is specified at the bottom. *Indicates a band remaining from previous staining, before stripping and reprobing the membranes.

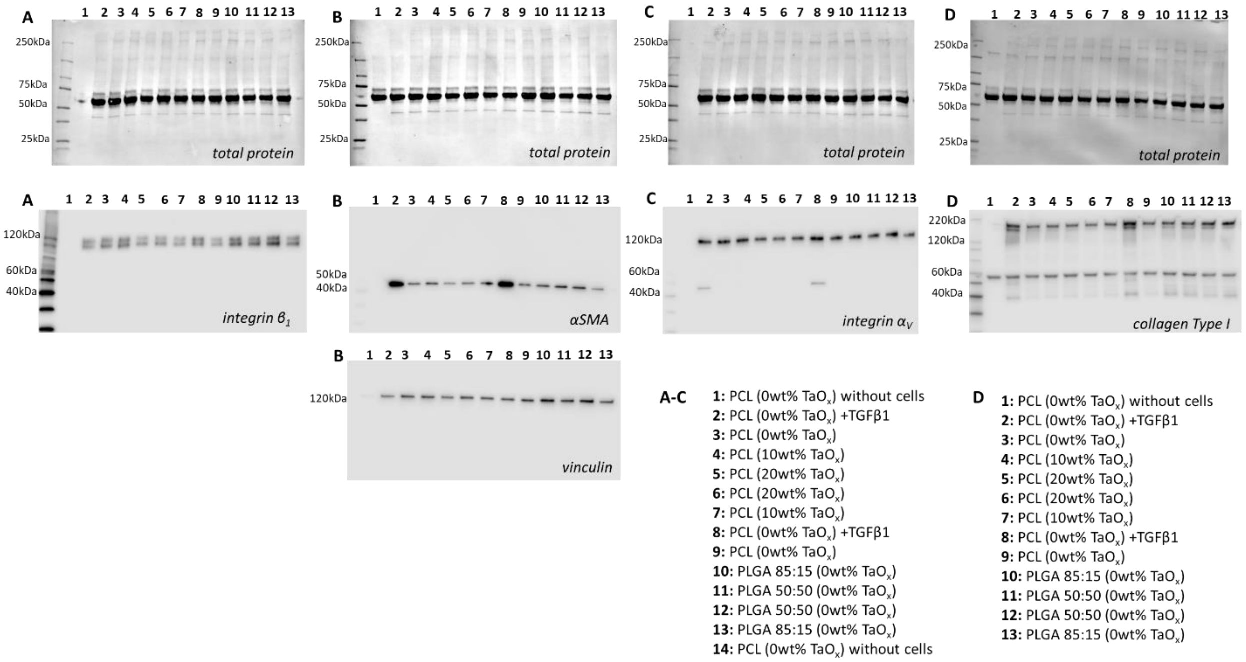


**Figure S4:** Western blots of porous devices at week 2 (biological replicate 2). Top row: total protein, second and third rows (A) integrin $\beta_1$, (B) αSMA, vinculin, (C) integrin $\alpha_V$, (D) collagen Type I. The sample loaded in each lane is specified at the bottom.

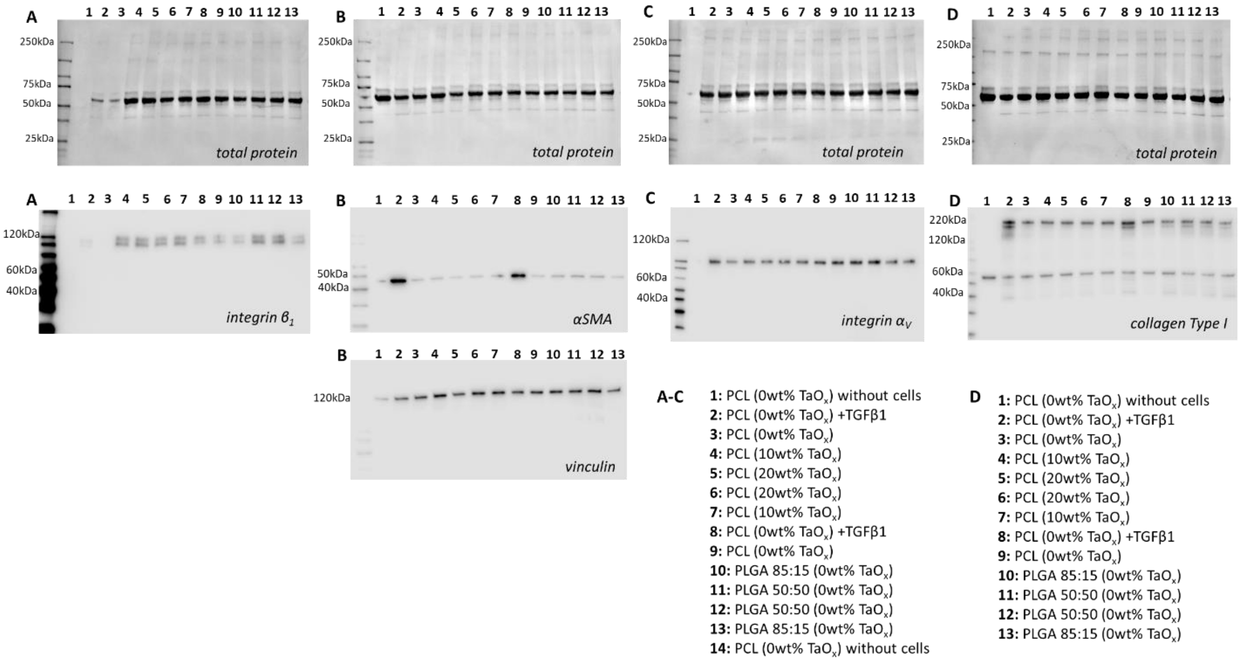


**Figure S5:** Western blots of porous devices at week 2 (biological replicate 3). Top row: total protein, second and third rows (A) integrin $\beta_1$, (B) αSMA, vinculin, (C) integrin $\alpha_V$, (D) collagen Type I. The sample loaded in each lane is specified at the bottom.

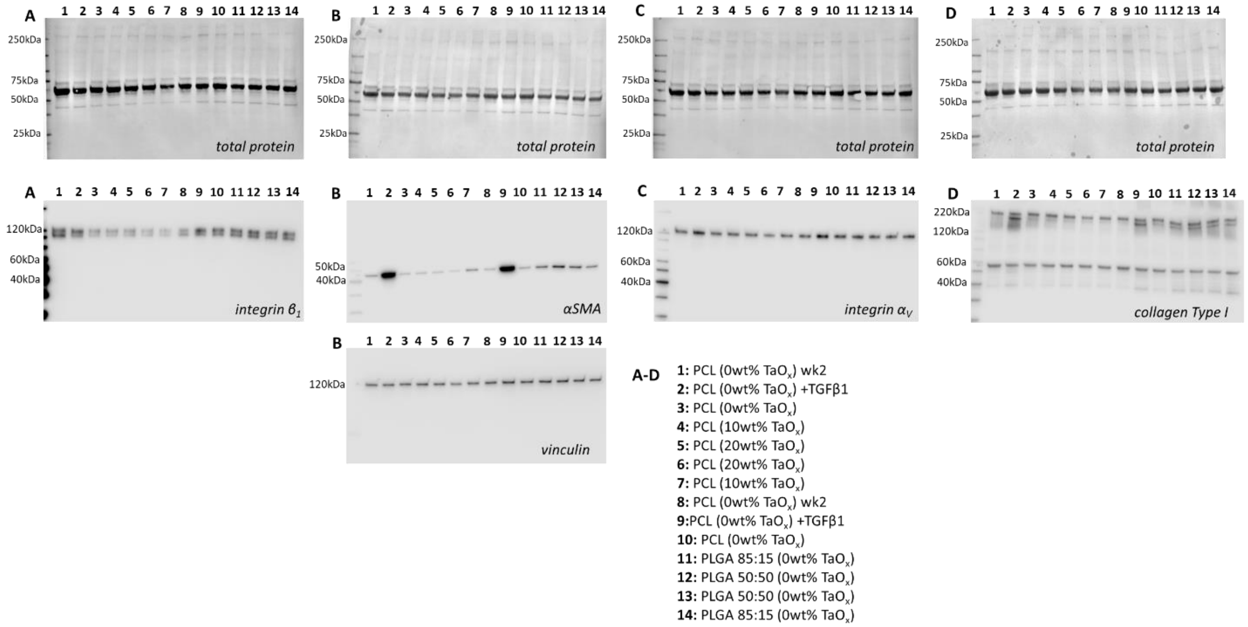


**Figure S6:** Western blots of porous devices at week 4 (biological replicate 1). Top row: total protein, second and third rows (A) integrin $\beta_1$, (B) $\alpha$SMA, vinculin, (C) integrin $\alpha_V$, (D) collagen Type I. The sample loaded in each lane is specified at the bottom.

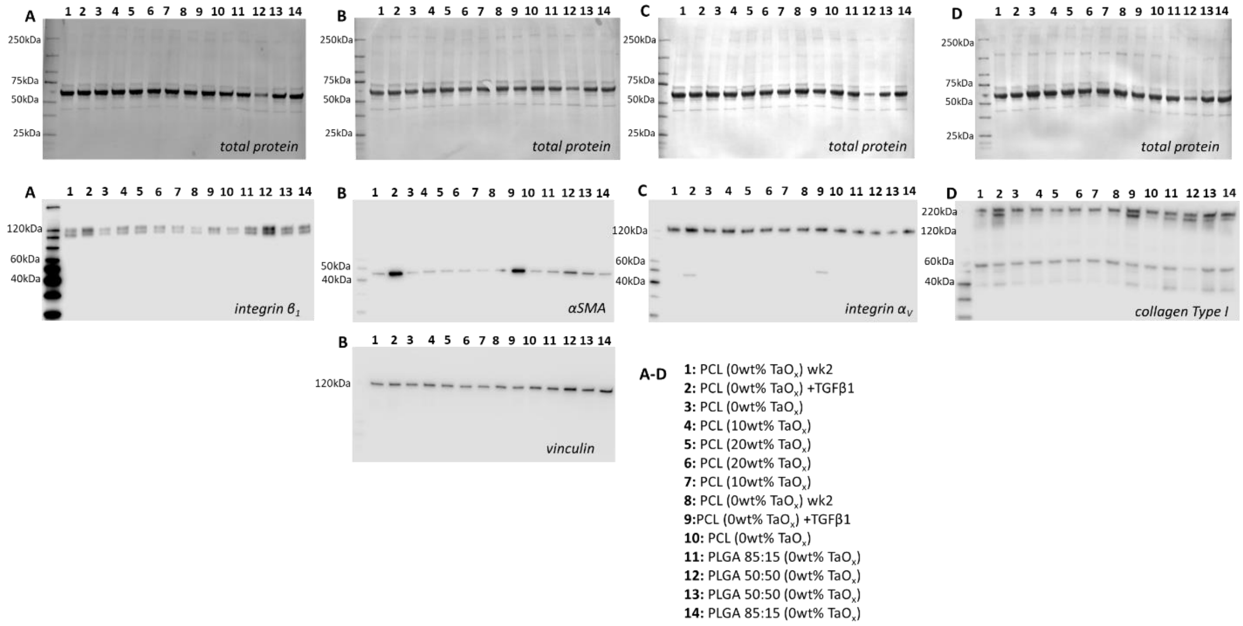


**Figure S7:** Western blots of porous devices at week 4 (biological replicate 2). Top row: total protein, second and third rows (A) integrin $\beta_1$, (B) $\alpha$SMA, vinculin, (C) integrin $\alpha_V$, (D) collagen Type I. The sample loaded in each lane is specified at the bottom.

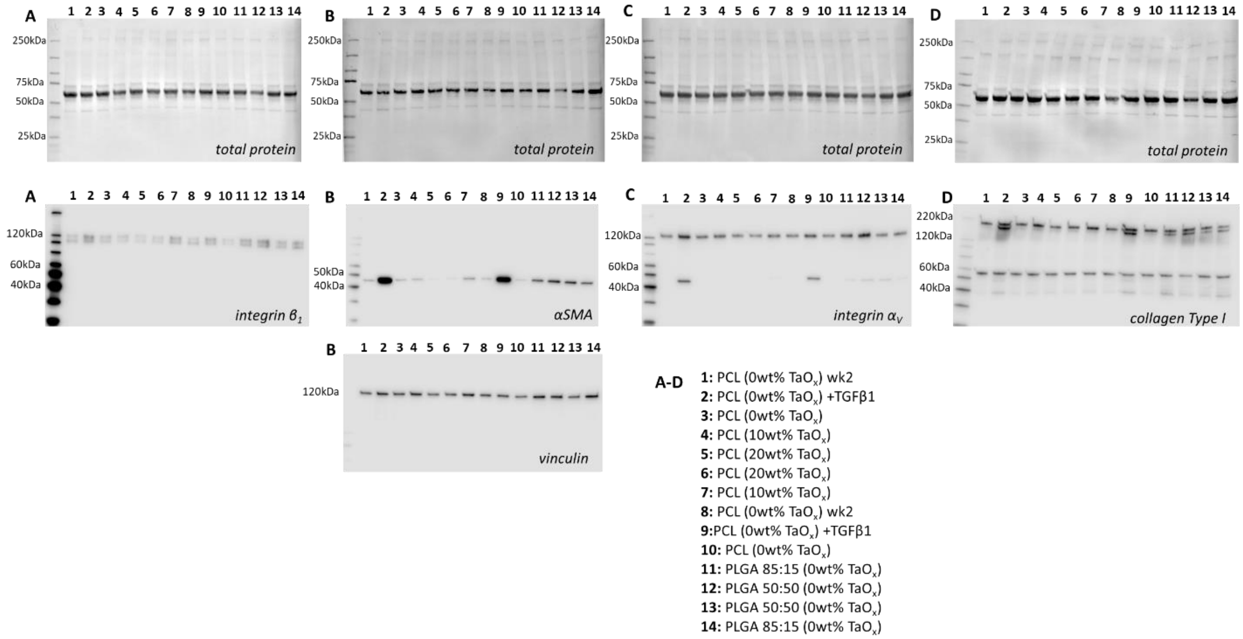


**Figure S8:** Western blots of porous devices at week 4 (biological replicate 3). Top row: total protein, second and third rows (A) integrin $\beta_1$, (B) αSMA, vinculin, (C) integrin $\alpha_V$, (D) collagen Type I. The sample loaded in each lane is specified at the bottom.

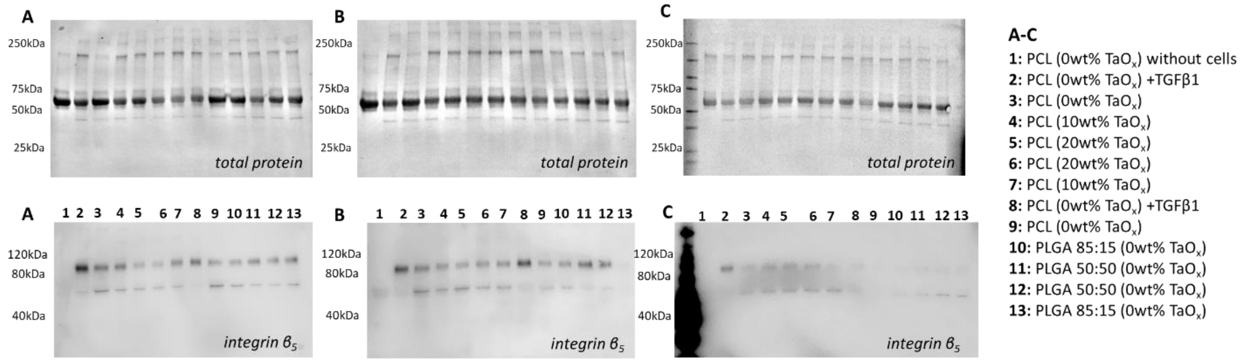


**Figure S9:** Western blots of porous devices at week 2: (A) biological replicate 1 (B) biological replicate 2 (C) biological replicate 3. Top row: total protein, second row: integrin $\beta_5$. The sample loaded in each lane is specified in the key. Protein signal was low compared to the protein ladder, so it was excluded in all blots except C.

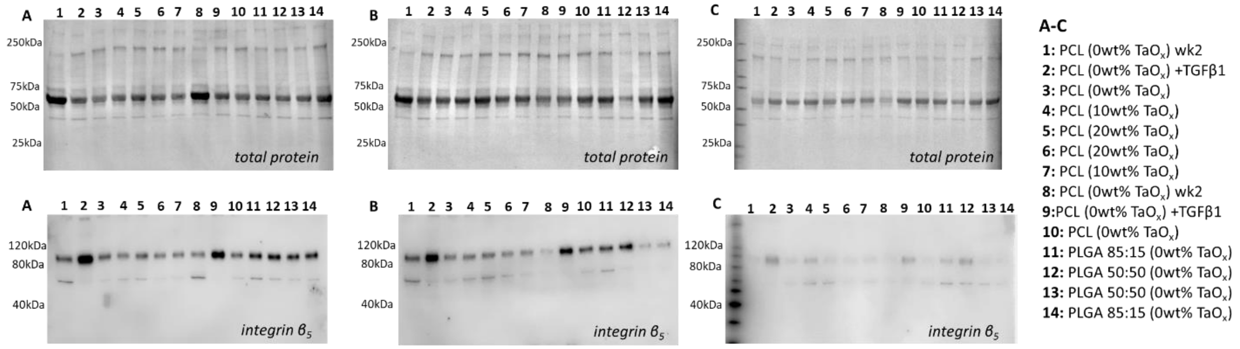


**Figure S10:** Western blots of porous devices at week 4: (A) biological replicate 1 (B) biological replicate 2 (C) biological replicate 3. Top row: total protein, second row: integrin $\beta_5$The sample loaded in each lane is specified in the key. Protein signal was low compared to the protein ladder, so it was excluded in all blots except C.

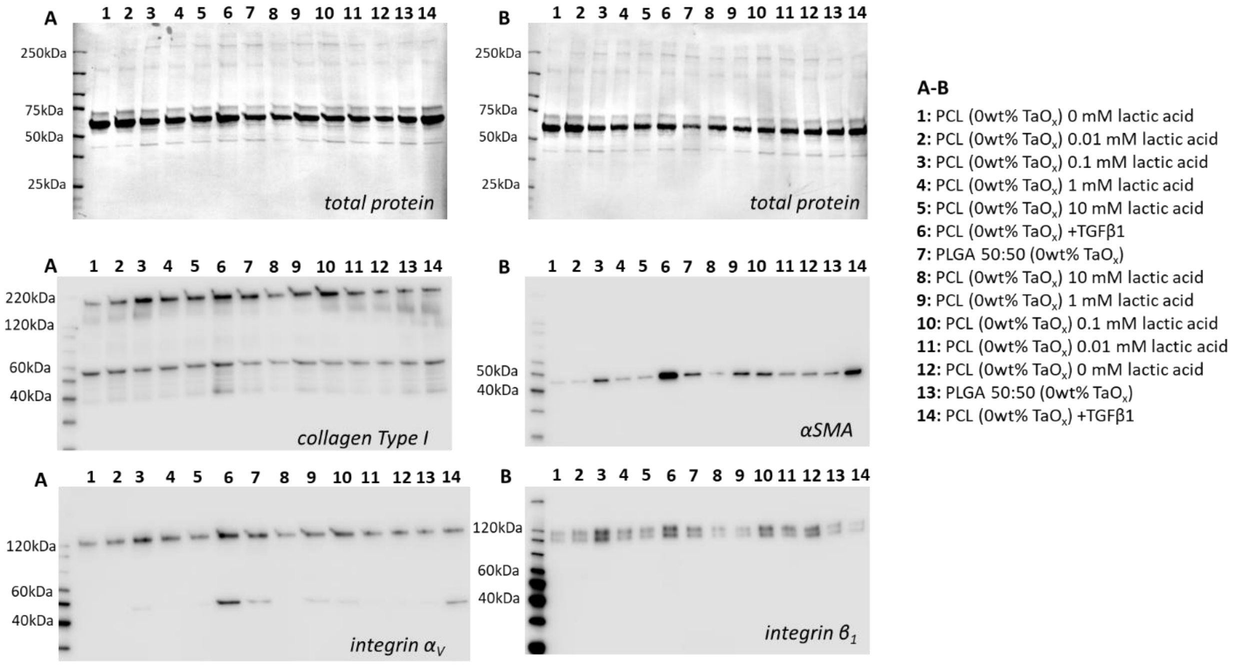


**Figure S11:** Western blots of films with lactic acid supplementation at week 2 (biological replicate 1). Top row: total protein, second row: (A) collagen Type I, integrin $\alpha_V$, (B) αSMA, integrin $\beta_1$. The sample loaded in each lane is specified in the key.

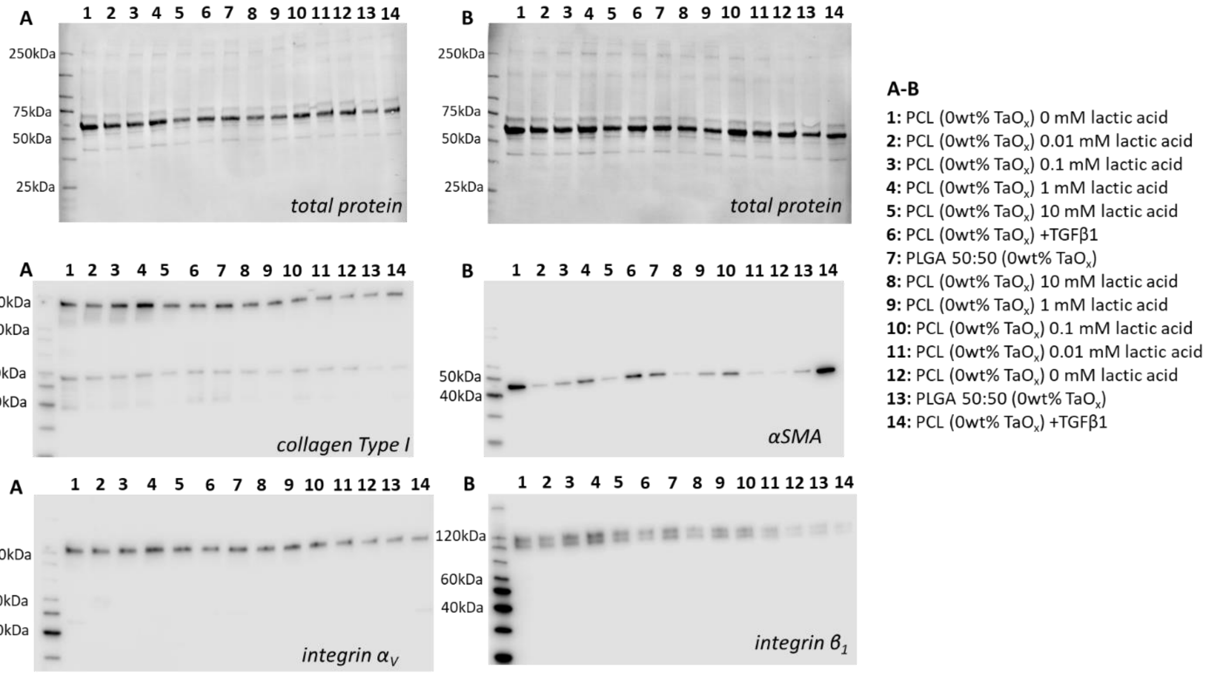


**Figure S12:** Western blots of films with lactic acid supplementation at week 2 (biological replicate 2). Top row: total protein, second row: (A) collagen Type I, integrin $\alpha_V$, (B) αSMA, integrin $\beta_1$. The sample loaded in each lane is specified in the key.

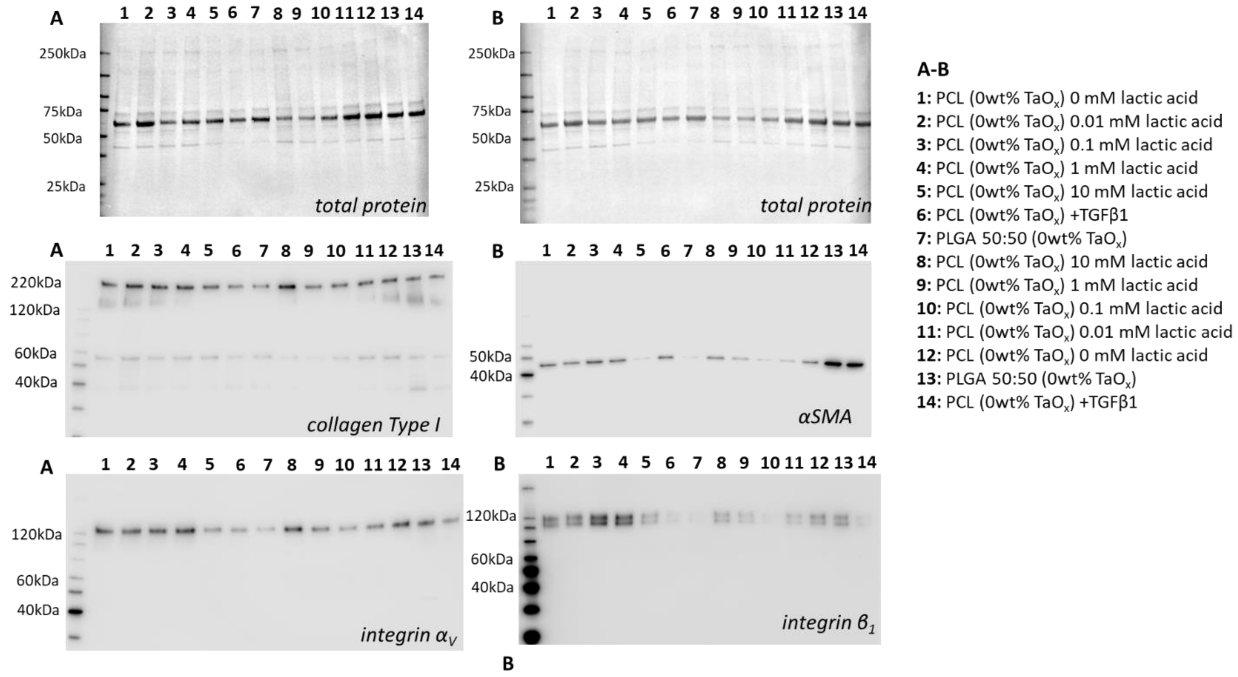


**Figure S13:** Western blots of films with lactic acid supplementation at week 2 (biological replicate 3). Top row: total protein, second row: (A) collagen Type I, integrin $\alpha_V$, (B) αSMA, integrin $\beta_1$. The sample loaded in each lane is specified in the key.
